# Serological evidence of dengue fever and associated factors in health facilities in Borena Zone, Southern Ethiopia

**DOI:** 10.1101/502617

**Authors:** Eshetu Nigussie Geleta

## Abstract

**Background:** Dengue fever is a re-emerging public health threat in Ethiopia. Yet, little is known about the epidemiology and risk factors. In this study the seroprevalence and associated risk factors of dengue virus infection were assessed in Borena Zone health facilities.

**Methods:** An institution based cross-sectional study was conducted from May to August, 2016. A total of 519 consecutive acute febrile patients attending the outpatient departments of Teltelle health Center, Yabello and Moyale Hospital were enrolled. Data on socio-demographic and environmental risk factors were collected using structured questionnaire. Three to five milliliter blood samples were collected from all participants and screened for dengue virus exposure using indirect immunofluorescent assay.

**Result:** The overall prevalence of anti-DENV IgG and IgM was 22.9% and 7.9% respectively. The relatively higher IgM versus IgG, absence of trend with age and little or no correlation with the assessed possible risk factors except being male (AOR=1.72; 95%CI 1.01-2.94), place of residence (AOR=0.37;95%CL 0.21-0.64) that had higher rate of exposure and recall of a recent mosquito bite (AOR=2.98; 95%CI 1.51-5.89) probably imply recent and/or ongoing active transmission.

**Conclusion:** This study showed dengue fever could potentially emerge as public health threat in the study area. On top, the observed low awareness of participants underline the urgent need for further systematic studies to determine the environmental, and host factors that determine the extent of exposure to dengue virus infection in the area for appropriate control and prevention planning.

**Author summary:** Dengue fever is a mosquito-borne viral disease of global health problem where *Aedes aegypti* mosquitoes are the main vector. It is endemic in most tropical and sub-tropical countries with an estimated 96 million infections resulting in clinical disease annually. It is unrecognized and underreported in Africa, particularly in Ethiopia. So, the current study were conducted among febrile patients who were attending health institutions to document seroprevalence and associated risk factors of DENV infection in the southern part of the country. The study stated the presence of antibodies against DENV infection in study areas, the ringing bell message for those who were involved in health sectors. Gender and residence were significantly associated with the prevalence of anti-DENV IgG seropositivity. In addition, individuals who have experience of recent mosquito bite were identified as the risk factors of DENV infection. Therefore, I recommend that preventive measures should be considered and nationwide surveillance should be carried out at nationwide.

## Introduction

Dengue fever (DF) is the most rapidly spreading mosquito-borne disease and the major public health problem on the world [1]. Dengue is a viral disease caused by dengue virus serotypes (DENV1-4) of the genus Flavivirus. Dengue virus is a non-segmented, positive-sense, singlestranded, enveloped RNA virus; transmitted primarily by the bite of *Aedes aegypti* and *Aedes albopictus* [2, 3]. Infections can also be transmitted through blood transfusion, organ transplantation and possibly vertically from mother to child [4, 5]. The virus distributed in more than 100 countries in tropical and subtropical areas; across the Americas, East Mediterranean, Western Pacific, Africa, South-East Asia and Europe [6]. The dengue infection has been reported in most African countries, especially in Eastern Africa including Ethiopia [3, 7, 8]. More than 390 million people are exposed to DENV each year resulting in 96 million annual cases of viral associated disease globally [9]. The World Health Organization (WHO) has reported 500,000 people develop severe disease each year, and about 1,250 die [10]. Although dengue has a global distribution, the majority of cases were from WHO South-East Asia region together with Western Pacific region bears nearly 75% of the global disease burden [11]. Recently, there were few dengue infection reports in Ethiopia, specifically eastern parts of the country Dire Dawa and Somali regions [3, 8, 12]. Several factors are related to the increase of dengue incidence in the Ethiopia. Among the most important ones are uncontrolled urbanization and absence of standardized public services such as water supply, sewage, and waste disposal [3].

Dengue virus infection produces a spectrum of clinical illness, ranging from an asymptomatic or mild febrile illness to classic DF to the most severe form of illness, dengue hemorrhagic fever (DHF) and dengue shock syndrome (DSS) [13]. DHF and DSS cases have also been increasingly recognized in South Asia, Latin America and the Pacific [14, 15], with pediatric cases being more common. Also DF and DHF/DSS have become more common in adults [16]. Dengue fever is clinically difficult to diagnose, especially in developing countries with no established dengue diagnostic techniques and could easily be mistaken for malaria, typhoid or unknown febrile illnesses [17]. The studies have reported that human antibody responses after dengue virus infection were highly cross-reactive with other arboviruses like Zika virus [18, 19].

Few studies of dengue infection have been carried out in Ethiopia; even though unknown causes of acute febrile illnesses are common. A confirmed DF case was reported for the first time in Ethiopia in Dire Dawa city in 2013 [8]. Later studies were conducted in somali region and northwestern parts of the country [3, 12]. However, data are not available on the DF in Southern Ethiopia. Thus, the aim of this study was to generate baseline data on the prevalence of DF and associated risk factors in acute febrile patients in health facilities with catchments from the Borena Zone. This study will be helpful in providing information on DENV infection to healthcare authorities for better clinical management of patients and to design and implement appropriate control measures.

## Materials and Method

### Ethical consideration

Ethical clearance was obtained from the Institutional review board of Hawassa University College of Medicine and Health Sciences, Oromia Regional Health Bureau Ethics Review Committee and AHRI/ALERT Ethics Review Committee. Before data collection, patients were informed about the objective and purpose of the study, about their right not to participate on the study or withdraw at any point in time. Personal privacy and dignity was respected. Data was collected after obtaining participants’/guardians’ informed written consent. Assent was also sought in cases the study participants were children under 18 years old. All samples and forms containing patient information had no name or information that can identify a particular participant; and data was analyzed and interpreted in aggregate.

### Study Site, design, period and patient’s characteristics

A health facility based cross-sectional study was carried out from May to August 2016 in Borena Zone: Yabello Hospital, Moyale Hospital and Teltelle Health center. A Zone is located in southern part of Ethiopia, bordering Kenya. The climate of the area is arid; mean annual rain fall of 400-700 mm in two rainy seasons (spring and autumn), and mean annual temperature ranging from 25-37°C. The study population consisted of all consecutive patients presenting with acute febrile illness at the outpatient departments during the study period. The sample size was 519.

### Data collection and Laboratory test

Data collectors interviewed the study participants using pretested structured questionnaire on socio-demographic and other risk factors such as use of bed net, trees around compound, use of mosquito repellent, presence of stagnant water around compound, and stay outside at night time. A 3-5 ml blood samples were collected, clotted and centrifuged at 1300r/minute. Separated sera were transported using liquid nitrogen (-170°c) to Hawassa University Referral Hospital, and stored in deep freezer (-80°c). Sera were transported using dry ice and screened at AHRI laboratory for DENV IgG and IgM using EUROIMMUN biochips indirect immunofluorescent assay (IIFA) kit (Medizinische Labordiagnostika AG-Germany) according to the manufacturer’s manual [20].

### Data analysis

Data was analyzed using SPSS version-20 window. Results were summarized using descriptive statistics and bivariate analysis. To control for possible effect of confounding, variables found to have an association with the outcome variable at P-value of 0.25 were entered into multivariable logistic regression model. Associations between independent and outcome variables were assessed and its strength was described using odds ratio with its 95% confidence intervals. P-value <0.05 was accepted as statistically significant.

## Results

### Socio-demographic characteristics

A total of 519 participants were investigated during the study period. Two hundred six (39.7%) of the study participants were from Teltelle health center, 36.6% were from Moyale hospital, and the remaining study participants (23.7%) were from Yabello hospital. The mean age of the participants was 25.5 years (range, 1 to 80 years, standard deviation 1.54), and those in the age range 15-29 years accounted 49.3%. Male participants accounted 51.3% with male to female ratio of 1:0.95. Substantial proportion of the study participants were rural residents (51.6%), illiterate (60.7%), and farmers by occupation (33.9%) (Table1).

**Table 1.**
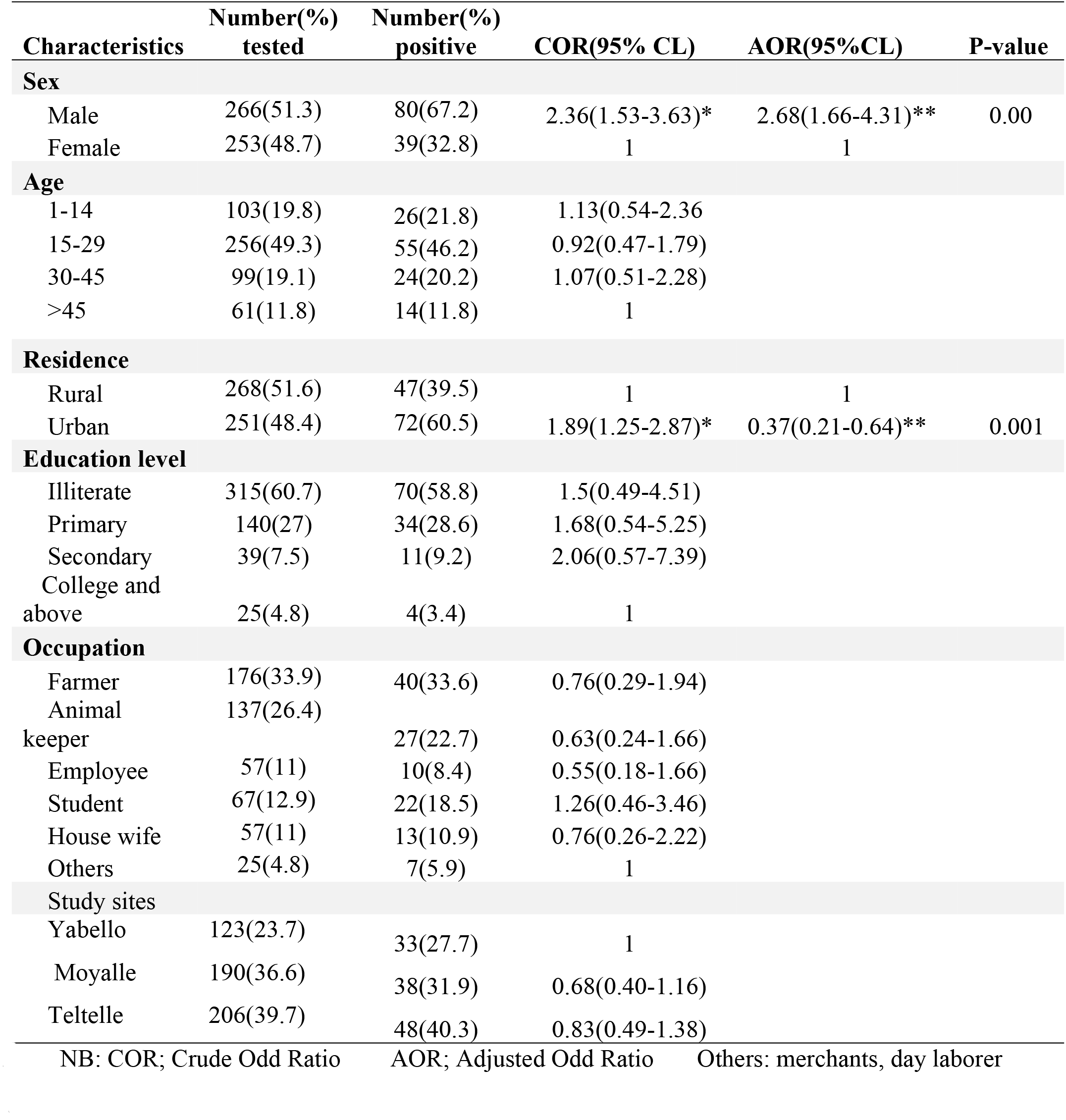
Dengue virus IgG seropositivity relation to socio-demographic characteristics of respondents attending health facilities in Borena Zone, Southern Ethiopia, 2016

### Seroprevalence of DENV infection

The overall prevalence of exposure to DF was found to be 22.9% and 7.9% for IgG and IgM respectively. Male participants (67.2%) had higher rate of DENV IgG compared to females (32.8%). With respect to age, the prevalence of DENV IgG was highest (46.2%) in age group 15-29 years and lowest (11.8%) in age group greater than 45 years. Further, the rate of DENV infection IgG exposure was higher among urban residents (60.5%), animal keeper (22.7%) and, illiterate individuals (58.8%), and Teltelle health center (40.3%) (Table1). Overall, 20 of 119 (48.8%) age group 15-29 years and 11 of 119 (26.8%) age group 1-14 years had IgM antibodies which suggest that recent infections with DENV were occurring (Table 3). In bivariate analysis, the association that yielded a p-value less than 0.25 with DENV IgG was gender and place of residence while age, study area, occupation and educational status of the study participants were not significantly associated. In multivariable logistic regression analysis male participants were at higher odds of having DENV IgG infection (AOR 2.68, 95% CI 1.66- 4.31) compared to females. Those study participants who lived in urban areas were 0.37 times (AOR = 0.37; 95% 0. 21-0.64) more likely to have anti-DENV IgG seropositivity than those who lived in rural areas (Table 1). The prevalence of DENV-3 IgG was highest (19.8%) and DENV-1 was lowest (8.3). Regarding IgM, both DENV-2 and DENV-3 were equally the highest (Table 2).

**Table 2.** Distribution of Dengue infection antibodies by serotypes.

|  | **IgG** | **IgM** |
| --- | --- | --- |
| **Dengue virus serotypes** | **Pos No (%)** | **Pos No (%)** |
| DENV-1 | 43(8.3) | 28(5.4) |
| DENV-2 | 71(13.7) | 34(6.6) |
| DENV-3 | 103(19.8) | 34(6.6) |
| DENV-4 | 66(12.7) | 31(6) |

**Table 3.** Dengue virus IgM seropositivity relation to socio-demographic characteristics of respondents attending health facilities in Borena Zone, Southern Ethiopia, 2016

| Characteristics | Number(%)<br>tested | Number(%)<br>positive | COR(95% CL) | AOR(95%CL) | P-value |
| --- | --- | --- | --- | --- | --- |
| **Sex** |  |  |  |  |  |
| Male | 266(51.3) | 16(39) | 1 |  |  |
| Female | 253(48.7) | 25(61) | 1.71(0.89-3.29) |  |  |
| **Age** |  |  |  |  |  |
| 1-14 | 103(19.8) | 11(26.8) | 2.31(0.62-8.64) |  |  |
| 15-29 | 256(49.3) | 20(48.8) | 1.64(0.47-5.70) |  |  |
| 30-45 | 99(19.1) | 7(17.1) | 1.47(0.36-5.92) |  |  |
| >45 | 61(11.8) | 3(7.3) | 1 |  |  |
| **Residence** |  |  |  |  |  |
| Rural | 268(51.6) | 16(39) | 1 |  |  |
| Urban | 251(48.4) | 25(61) | 1.74(0.91-3.35) |  |  |
| **Education level** |  |  |  |  |  |
| Illiterate | 315(60.7) | 20(48.8) | 0.49(0.14-1.80) |  |  |
| Primary | 140(27) | 16(39) | 0.95(0.25-3.52) |  |  |
| Secondary | 39(7.5) | 2(4.9) | 0.39(0.06-2.56) |  |  |
| College and above | 25(4.8) | 3(7.3) | 1 |  |  |
| **Occupation** |  |  |  |  |  |
| Farmer | 176(33.9) | 10(24.4) | 1.45(0.18-11.80) |  |  |
| Animal keeper | 137(26.4) | 8(19.5) | 1.49(0.18-12.45) |  |  |
| Employee | 57(11) | 5(12.2) | 2.31(0.26-20.84) |  |  |
| Student | 67(12.9) | 10(24.4) | 4.21(0.51-34.74) |  |  |
| House wife | 57(11) | 7(17.1) | 3.36(0.39-28.87) |  |  |
| Others | 25(4.8) | 1(2.4) | 1 |  |  |
| **Study sites** |  |  |  |  |  |
| Yabello | 123(23.7) | 9(22) | 1 |  |  |
| Moyalle | 190(36.6) | 14(34.1) | 1.01(0.42-2.41) |  |  |
| Teltelle | 206(39.7) | 18(43.9) | 1.21(0.53-2.79) |  |  |

NB: COR; Crude Odd Ratio AOR; Adjusted Odd Ratio Others: merchants, day laborer

**Table 4.** Dengue virus IgG seropositivity relation to knowledge and environmental characteristics of respondents attending health facilities in Borena Zone, Southern Ethiopia, 2016

| Characteristics | Number(%)<br>tested | Number(%)<br>positive | COR (95% CL) | AOR(95% CL) | P-value |
| --- | --- | --- | --- | --- | --- |
| **Heard about DF** |  |  |  |  |  |
| Yes | 198(38.2) | 52(43.7) | 1 |  |  |
| No | 321(61.8) | 67(56.3) | 0.74(0.49-1.12) |  |  |
| **Mode of transmission** |  |  |  |  |  |
| Mosquito | 50(9.6) | 16(13.4) | 1 |  |  |
| By blood | 14(2.7) | 5(4.2) | 1.18(0.34-4.09) |  |  |
| Do not know | 455(87.8) | 98(82.4) | 0.58(0.31-1.10) |  |  |
| **Stagnant water** |  |  |  |  |  |
| Yes | 162(31.2) | 36(30.3) | 1 |  |  |
| No | 357(68.8) | 83(69.7) | 1.06(0.68-1.65) |  |  |
| **Trees around compound** |  |  |  |  |  |
| Yes | 333(64.2) | 80(67.2) | 1.19(0.77-1.84) |  |  |
| No | 186(35.8) | 39(32.8) | 1 |  |  |
| **Stay outside at night** |  |  |  |  |  |
| Yes | 248(47.8) | 62(52.1) | 1.25(0.83-1.88) |  |  |
| No | 271(52.2) | 57(47.9) | 1 |  |  |
| **Recent mosquito bite** |  |  |  |  |  |
| Yes | 300(57.8) | 84(70.6) | 2.04(1.32-3.18)* | 2.23(1.39-3.56)** | 0.001 |
| No | 219(42.2) | 35(29.4) | 1 | 1 |  |
| **Bed net use** |  |  |  |  |  |
| Yes | 333(63.6) | 81(68.1) | 1 |  |  |
| No | 186(36.4) | 38(31.9) | 1.25(0.81-1.94) |  |  |
| **Repellent use** |  |  |  |  |  |
| Yes | 21(4.1) | 6(5) | 1 |  |  |
| No | 498(95.9) | 113(95) | 0.73(0.28-1.94) |  |  |

NB: COR; Crude Odd Ratio AOR; Adjusted Odd Ratio Others: merchants, day laborer

Regarding the general awareness about DF, 38.2% of the participants had heard about this virus infection, and 9.6% responded DENV is transmitted by mosquito. Respondents were asked about the environmental exposures associated with mosquito-borne illnesses in their dwelling areas. Those who reported the existence of stagnant water and trees nearby their dwelling were 31.2% and 64.2%, respectively. Above 57% of respondents reported recent mosquito bites while they stayed outside during night time (47.8%). Three hundred thirty three (63.6%) of the study participants reported they slept under mosquito nets; of which 20.2 and 41.4% used bed net always and sometimes, respectively. However, only 4.1% used mosquito repellents on day or night time (Table 4).

The seropositivity of DENV IgG was 43.7% in those who heard about the virus and 13.4% in those who responded mosquito transmits the infection. The rate of exposure was observed 52.1% and 68.1% among those who responded they had a habit of staying outside during night and used bed net. A recent experience of having a mosquito bite (70.6%) was the only factor that significantly influenced the rate of DENV IgG exposure in bivariate analysis. However, use of mosquito repellant, awareness of DF, knowledge of transmission route, presence of tree around compound, habit of staying out side home in night time and a use of bed net were not significantly associated (p-value > 0.05). The association between recent mosquito bite and DENV IgG infection was found to be statistically significant in a multivariable logistic regression analysis (AOR=2.23; 95%CI, 1.39-3.56, p=0.001) (Table 4).

## Discussion

Recently, dengue fever infection has been considered an emerging public health problem in several African countries with risk of severe infections [21, 22]. Most febrile cases are routinely diagnosed and treated for typhoid and/or malaria without proper investigation for other conditions including viral infections. In Ethiopia where various mosquito-borne diseases are common, little is known about the epidemiology of arboviruses including DENV. However, the 2013 and 2014 DF outbreaks in Dire Dawa and Godey [8, 12] that caused many morbidities and mortalities calls for systematic investigations to better describe the epidemiology of DF in various localities. Specially in relation to a worsening situation in climate change, which supports the emergence and re-emergence of vector-borne diseases, the need to have a strong surveillance system is critically important. This study assessed the prevalence of DENV infection and its associated risk factors in health facilities in Borena Zone where febrile illness is common.

The seroprevalence of exposure to DENV IgG among febrile patients in the study area were 22.9%. This result is in agreement with findings reported in Djibouti, 21.8% [23] and in Northern Province of Sudan, 24% [24]. However, the observed rate of DENV exposure was lower than results in Dire Dawa, 56.8% [8], northern Ethiopia, 33.3% [3], Eritrea, 33.3%[25], Kassala, Eastern Sudan, 71.7% [26], and El Gadarif state, Sudan, 47.6% [27]. In contrast, the prevalence of anti-DENV IgG exposure in this study was higher compared to the rates 7.7% and 12.5% in Tanzania and Kenya respectively [28, 29]. These discrepancies may be due to difference in distribution of risk factors and variable climatic conditions by geographical regions, diversity of the studied populations, and difference in diagnostic performance of the employed laboratory methods. For example, some studies analyzed samples using laboratory techniques such as ELISA, PRNT and PCR which are more sensitive and specific compared to IIFA technique used in the current study. The high circulation of DENV in the study area could be attributed to several factors including misdiagnosis of febrile cases, the movement of migrants from endemic countries and the proliferation of breeding sites of *Aedes mosquitoes*. And also one-fourth of the study participants had antibody against DENV infection, dengue was under recognized and underreported in Ethiopia, which is in line with an earlier report in Africa [7]. The overall prevalence of anti-DENV IgM seropositivity was 7.9% which indicates recent infection with DENV. Since IgM against the DENV infection can be usually detected after the first 5-7 days of infection [30, 31]. However, the possibility that as the IgM antibodies remain negative for the first few days, and also the IgM reactivity was non-specific; thus there is cross-reactive due to infection with another flavivirus [32].

This study showed that gender significantly influenced the rate of anti-DENV IgG exposure status where male participants were disproportionally infected, which is in agreement with the study conducted elsewhere [3, 33]. It might be due to that males are more likely to work in outdoor forested areas where they come into contact with vectors for DENV. In this study, those individuals who were dwelling in urban areas were more affected than those in the rural areas. This is in agreement with the studies elsewhere [3, 34, 35]. It was previously reported that the seropositivity rate for DENV, which is carried by common vectors *Ae. aegypti* and *Ae. albopictus*, was higher in the geographically central sites (urban centers) than villages [36]. Recent mosquito bite was significantly associated with anti-DENV IgG seropositivity. This is in line with the fact that mosquito bite exposes individuals to DF, and it may be the main mode of transmission in the study area. However, factors such as age, study site, occupation and educational status have little significance in influencing the rate of exposure to DENV in the current study.

Although this is the first study of seroprevalence and risk factors associated with DENV infection in Southern Ethiopia, the study has several limitations. IIFA was shown to have good performance as compared PRNT; its inherent cross reactivity to other Flaviviruses could not be ruled out. Moreover, no febrile community controls or convalescent sera, and as any health institution based study that used consecutive volunteering cases only the risk of introducing bias is unavoidable. Thus, the findings of this study may not be generalized to the population in the study area.

In conclusion, this study showed low awareness among participants and the potential that DENV could likely be public health significance in the study area. Thus, we recommend a community based survey in the study and adjacent communities to verify our findings and take appropriate public health measures to provide potential outbreaks. Further systematic studies should be conducted to determine the environmental, and host factors that determine the extent of exposure to DENV infection in the area for appropriate control and prevention planning.

**Abbreviation**
AHRI: Armauer Hansen research institute.
ELISA: Enzyme Immunosorbent assay
IIFA: Indirect Immunofluorescent Assay
IgG: Immunoglobulin G
IgM: Immunoglobulin M
PRNT: Plague Reduction Neutralization test
DF: Dengue fever
DENV: Dengue virus

## Acknowledgement

I thank the Oromia Regional Health Bureau, health Bureaus and the responsible officials and professionals in the study area for their cooperation. I am also most grateful to the study participants for volunteering.

## Supporting information

S1 STROBE Checklist. (DOC)

## References

1. Yong YK, Thayan R, Chong HT, Tan CT, Sekaran SD. Rapid detection and serotyping of dengue virus by multiplex RT-PCR and real-time SYBR green RT-PCR. Singapore medical journal. 2007 Jul;48(7):662.

2. Dhar-Chowdhury P, Paul KK, Haque CE, Hossain S, Lindsay LR, Dibernardo A, et al. Dengue seroprevalence, seroconversion and risk factors in Dhaka, Bangladesh. PLoS neglected tropical diseases. 2017 Mar 23;11(3):e0005475.

3. Ferede G, Tiruneh M, Abate E, Wondimeneh Y, Damtie D, Gadisa E, et al. A serologic study of dengue in northwest Ethiopia: Suggesting preventive and control measures. PLoS neglected tropical diseases. 2018 May 31;12(5):e0006430.

4. Stramer SL. The potential threat to blood transfusion safety of emerging infectious disease agents. Clinical Advances Hematology and Oncology. 2015; 13:420–2.

5. Weerakkody RM, Palangasinghe DR, Dalpatadu KP, Rankothkumbura JP, Cassim MR, Karunanayake P. Dengue fever in a liver-transplanted patient: a case report. Journal of medical case reports. 2014;8(1):378.

6. Kyle J, Harris E. Global spread and persistence of dengue. Annu Rev Microbiol. 2008; 62:71–92. https://doi.org/10.1146/annurev.micro.62.081307.163005 PMID: 18429680

7. Ananda A, Joel N, Kuritsky G, William L, Harold S, Margolis H. Dengue virus infection in Africa. Emerg Infect Dis. 2011; 17:1349–54 https://doi.org/10.3201/eid1708.101515 PMID: 21801609

8. Abyot BW, Mesfin M, Wubayehu K, Esayas K, Milliyon W, Abiy G, et al. The first acute febrile illness investigation associated with dengue fever in Ethiopia, 2013: A descriptive analysis. Ethiop J Health Dev. 2014; 28:155–61.

9. Guzman MG, Harris E (2015) Dengue. The Lancet 385: 453–465.

10. World Health Organization (WHO) Global strategy for dengue prevention and control 2012-2020. Geneva, Switzerland: World Health Organization, 2012.

11. WHO-TDR. Dengue: guidelines for diagnosis, treatment, prevention, and control-New Edition. Geneva, Switzerland. World Health Organization. 2009

12. Yusuf MA, Ali AS. Epidemiology of Dengue Fever in Ethiopian Somali Region: Retrospective Health Facility-Based Study. CAJPH. 2016; 2:51–6.

13. Martina BE, Koraka P, Osterhaus A (2009). Dengue virus pathogenesis, an integrated view. Clin. Microbiol. Rev., 22: 564–581.

14. Gupta N, Srivastava S, Jain A, Chaturvedi UC. Dengue in India. Indian J Med Res 2012; 136:373–390.PMID:23041731

15. Humayoun MA, Waseem T, Jawa AA, Hashmi MS, Akram J. Multiple dengue serotypes and high frequency of dengue hemorrhagic fever at two tertiary care hospitals in Lahore during the 2008 dengue virus outbreak in Punjab, Pakistan. International Journal of Infectious Diseases. 2010;14:e54–9.

16. Teixeira MG, Costa MC, Coelho G, Barreto ML. Recent shift in age pattern of dengue hemorrhagic fever, Brazil. Emerging infectious diseases. 2008;14(10):1663–.

17. Baba M, Saron MF, Vorndam A, Adeniji J, Diop O, Olaleye D. Dengue virus infections in patients suspected of malaria/typhoid in Nigeria. J Am Sci. 2009;5(5):129–134.

18. Priyamvada L, Quicke KM, Hudson WH, Onlamoon N, Sewatanon J, Edupuganti S, et al. Human antibody responses after dengue virus infection are highly cross-reactive to Zika virus. Proc Natl Acad Sci U S A. 2016;113:7852–7.

19. Dejnirattisai W, Supasa P, Wongwiwat W, Rouvinski A, Barba-Spaeth G, Duangchinda T,etal. Dengue virus sero-cross-reactivity drives antibody dependent enhancement of infection with zika virus. Nat Immunol. 2016;17:1102–8.

20. EUROIMMUN AG. Biochips mosaics and profile for detection of Flavivirus infections instructions for indirect immunoflourescent test, 2013.

21. Amarasinghe A, Kuritsk JN, Letson GW, Margolis HS. Dengue virus infection in Africa. Emerg Infect Dis. 2011;17(8):1349–1354.

22. Shepard DS, Undurraga EA, Halasa YA. Economic and disease burden of dengue in Southeast Asia. PLoS Negl Trop Dis. 2013;7(2): e2055

23. Andayi F, Charrel R, Kieffer A, Richet H, Pastorino B, Leparc I. A Sero-epidemiological Study of Arboviral Fevers in Djibouti, Horn of Africa. PLoS Negl Trop Dis. 2014; 8:e3299. https://doi.org/10.1371/journal.pntd.0003299 PMID: 2550269

24. Watts D, EI-Tigani A, Botros B, Salib A, Olmn J, McCarthy M. Arthropod-borne viral infections associated with a fever outbreak in the northern province of Sudan. J Trop Med Hyg. 1994; 97:228–30. PMID: 8064945

25. Abdulmumini U, Jacob D, Diana R, Araia B, Yohannes G, Goitom M, et al. Dengue fever outbreaks in Eritrea, 2005-2015. A case for strengthening surveillance, control, and reporting. Glob Health Res Policy. 2016; 1:17. https://doi.org/10.1186/s41256-016-0016-5 PMID: 29202065

26. Tajeldin A, AbdelAziem A, Mubarak K, Ishag A. Epidemiology of Dengue Infections in Kassala, Eastern Sudan. J Med Virol. 2012; 84:500–3. https://doi.org/10.1002/jmv.23218 PMID: 22246838

27. Eldigail MH, Adam GK, Babiker RA, Khalid F, Adam IA, Omer OH, Ahmed ME, Birair SL, Haroun EM, AbuAisha H, Karrar AE. Prevalence of dengue fever virus antibodies and associated risk factors among residents of El-Gadarif state, Sudan. BMC public health. 2018 Dec;18(1):921.

28. Francesco V, Emanuele N, Silvia M, Monica S, Maria G, Nazario B, et al. Seroprevalence of dengue infection: a cross-sectional survey in mainland Tanzania and on Pemba Island, Zanzibar. Int J Infect Dis. 2012; 16:e44–e6. https://doi.org/10.1016/j.ijid.2011.09.018 PMID: 22088862

29. Caroline O, Petronella A, Aymond N, Stella G, Cyrus W. Seroprevalence of Infections with Dengue, Rift Valley Fever and Chikungunya Viruses in Kenya. PLoS One. 2015; 10:e0132645. https://doi.org/10.1371/journal.pone.0132645 PMID: 26177451

30. Maria G, Scott B, Harvey A, Philippe B, Jeremy F, Duane J, et al. Dengue: a continuing global threat. Nat Rev Microbiol. 2010; 8:S7–S16. https://doi.org/10.1038/nrmicro2460 PMID: 21079655

31. Madara AA, Abdulraheem NO. Relative abundance of adult mosquitoes in University of Abuja Main Campus, Abuja FCT, Nigeria. Nig J Parasitol. 2013;34(2):1–5.

32. Calisher C, Karabatsos N, Dalrymple J, Shope R, Porterfield J, Westaway E, et al. Antigenic relationships between flaviviruses as determined by cross-neutralization tests with polyclonal antisera. J Gen Virol. 1989; 70:37–43.

33. Yik W, Tun Y, Li W, Lee C, Grace Y, Lyn J, et al. Seroepidemiology of Dengue Virus Infection Among Adults in Singapore. Ann Acad Med Singapore. 2009; 38:667–75. PMID: 19736569

34. Mohammed S, Syed M, Omrana P, Syed A, Mubarak K, Mutasim E, et al. Dengue fever in a border state between Sudan and Republic of South Sudan: Epidemiological Perspectives. J Public Health Epidemiol. 2013; 5:319–24. 56.

35. Sadia N, Muhammad A, Muhammad A, Ahmad R, Bahar M. The epidemiology of dengue fever in district Faisalabad, Pakistan. Int J Sci Res. 2015; 5

36. Gubler D. Dengue, Urbanization and Globalization: The Unholy Trinity of the 21(st) Century. Trop Med Health. 2011; 39:3–11.

